# Multiple roles of Pax6 in corneal limbal epithelial cells and maturing epithelial cell adhesion

**DOI:** 10.1101/2022.05.23.493090

**Authors:** Sweetu Susan Sunny, Jitka Lachova, Naoko Dupacova, Zbynek Kozmik

## Abstract

Mammalian corneal development is a multistep process, including formation of corneal epithelium (CE), endothelium and stroma during embryogenesis followed by postnatal stratification of the epithelial layers, and continuous renewal of the epithelium to replace the most outer corneal cells. Herein we employed Cre-*loxP* system to conditionally deplete Pax6 proteins in two domains of ocular cells, including the ocular surface epithelium (cornea, limbus and conjunctiva) or postnatal CE, via *K14-cre* or *Aldh3-cre*, respectively. Earlier and broader inactivation of Pax6 in the OSE resulted in thickened OSE with CE and limbal cells adopting the conjunctival keratin expression pattern. More restricted depletion of Pax6 in postnatal CE resulted in the abnormal cornea marked by reduced epithelial thickness despite of increased epithelial cell proliferation. Immunofluorescence studies showed loss of Keratin 12, an intermediate filament and diffused expression of adherens junction components, together with reduced tight junction protein, Zonula occludens-1 (ZO-1). Furthermore, expression of Keratin 14, basal cell marker in apical layers, indicates impaired differentiation of corneal epithelial cells. Collectively, our data demonstrate that Pax6 is essential for maintaining proper differentiation and strong intercellular adhesion in postnatal corneal epithelial cells, whereas limbal Pax6 is required for preventing the outgrowth of conjunctival cells to the cornea.

## Introduction

The complex process of vision starts with the light transmission and refraction through the multi-layered cornea, a transparent avascular tissue, which contributes to 60-70 % of the eye’s total refractive power as well functions as a protective barrier [1]. The cornea comprises mainly of three compartments; outermost non-keratinised stratified squamous epithelium, central collagenous stroma with sparsely dispersed keratocytes and innermost monolayered endothelium [2, 3]. Any defects in corneal development or maintenance of its homeostasis results in loss of vision [4, 5].

Mammalian corneal development is a multistep process originating at the early optic cup stage with the formation of the presumptive corneal epithelium from the surface ectoderm, followed by formation of the corneal endothelium and keratinocytes during two migratory waves of the neural crest-derived periocular mesenchyme. Postnatal stratification of the epithelial layers, and continuous renewal of the epithelium are central processes of adult corneal cell biology [6–10]. Mouse corneal development is a leading mammalian model to understand cellular, molecular and genetic mechanisms underlying corneal development and homeostasis. Coincident with eyelid opening around postnatal day14 (PN14), 1-2 layered corneal epithelium (CE) divides and differentiate into 4-6 layers by PN21 and form mature 6-8 layered tissue by 8-10 weeks after birth [2, 3, 11]. In the mature cornea, most superficial cells are shed off and replaced regularly by differentiated cells that move apically from the underlying layers, which are replenished by centripetal migration of stem cells from the limbus located at the juncture between cornea and conjunctiva [11–15].

A paired domain and homeodomain-containing transcription factor Pax6 is an evolutionarily conserved regulator of eye development in both vertebrates and invertebrates [16, 17].

In the mouse, Pax6 is expressed in the surface ectoderm that gives rise first to the lens prospective ectoderm and prospective corneal epithelium [18]. Pax6 is also expressed in the corneal limbus where stem cells reside as well as in the conjunctiva [19, 20].

A number of mutated *Pax6* alleles have been used to dissect both cell type- and stage-specific roles of this transcription factor in the mouse eye development, including the cornea [21–33]. Heterozygous mutations in both human *PAX6* and mouse *Pax6* genes cause a range of ocular and postnatal developmental abnormalities (i.e. human aniridia and Peters’ anomaly) that affect all ocular tissues at variable degrees depending on the type and location of each individual mutation [34, 35]. In cornea, the prevailing PAX6-associated defects are aniridia-associated keratopathy [36, 37] and limbal stem cell deficiency [38]. In contrast, *Pax6* homozygous mutations such as *Sey/Sey* lack the eyes, exhibit major brain, olfactory placode and pancreas abnormalities and die at birth [39].

Prenatally, *Pax6^+/-^* corneas exhibit thin epithelium and thickened hypocellular stroma. Postnatally, cornea collapses further with inflammatory cell infiltration, neovascularisation, erosions, and goblet cells accumulation in the epithelium, leading to corneal opacity [24–27, 40]. Further studies on factors contributing to corneal pathogenesis in *Pax6^+/-^* mutants revealed enhanced cell turnover, delayed expression of cytokeratin 12 and defective centripetal migration of limbal stem cells (LSC) [24–26, 41]. However, the lethality of homozygous Sey mutants, the consecutive induction of lens and cornea from SE, the impact of the lens in morphogenesis of anterior segment [22, 23, 42, 43] and continued expression of Pax6 in the adult lens and cornea [44], makes it challenging to dissect both spatial and temporal functions of Pax6 in CE using classical knockout models.

Analysis of mouse *Pax6* heterozygous mutations offer some insight into its corneal function; however, loss-of-function studies resulting in complete depletion of Pax6 protein provides key mechanistic insights how this transcription factor functions in different corneal cellular compartments. Here we thus took advantage of the Cre-*loxP* system for selectively inactivating Pax6 either in the ocular surface epithelium (OSE) (cornea, limbus, and conjunctiva) or CE. We used two different Cre drivers: *K14-Cre*, where cre activity is controlled by keratin K14 promoter, specific for basal keratinocytes in OSE [45] and *Aldh3-Cre*, where cre activity is controlled by Aldh3/Aldh1a3 promoter-specific for postnatal CE [46]. Pax6 ablation using *K14-Cre* from OSE at early postnatal stages (PN1-PN9) resulted in conjunctivalisation of the CE and limbal epithelium, suggesting the importance of limbal Pax6 in preventing the overgrowth of conjunctival epithelia to cornea. In contrast, specific deletion of Pax6 from CE upon eyelid opening leads to an abnormal thin cornea with defective cell-cell adhesion, thus providing direct evidence for the function of Pax6 in postnatal corneal development.

## Materials and Methods

### Ethics statement

The housing of mice and in vivo experiments were done in concurrence with the European Communities Council Directive 86/609/EEC of 24 November 1986 and institutional and national guidelines. All experimental methods and animal care were approved by the Animal Care Committee of the Institute of Molecular Genetics (no. 71/2014). This work did not include human subjects.

#### Mouse strains

Following genetically modified mouse strains were used for this study: *Aldh3-Cre* [46], *K14-Cre* [45] and *Pax6^fl/fl^* [32].

#### Histology and immunofluorescence

Eyeballs from pups of desired age were enucleated, fixed in 4% formaldehyde in 1x PBS (Phosphate Buffered Saline), for overnight at 4°C, then washed 3×10 minutes in 1x PBS, dehydrated in ethanol series, cleared in xylene, and then processed to paraffin wax.

For histological analysis, sections (6μm) were deparaffinised, rehydrated with ethanol series and stained with Haematoxylin and Eosin (H & E) and mounted in glycerol. For immunofluorescence, deparaffinised sections were heat-treated with sodium citrate buffer (10mM sodium citrate buffer, 0.05% Tween 20, pH-6.0) for antigen retrieval. Non-specific binding were blocked by incubation in 10% BSA (Bovine serum albumin) / 0.1% PBT (PBS with 0.1% Tween-20) for 30 minutes and then incubated with primary antibodies in 1%BSA / 0.1% PBT for overnight. Subsequently, samples were washed in 1x PBS, for 3×10 minutes, followed by incubation for one hour in secondary antibodies in 1% BSA / 0.1% PBT, washed in 1x PBS, 2×10 minutes, counterstained with DAPI(1μg/ml) and mounted using Mowiol (Sigma).

For staining with antibodies related to adhesion, eyeballs collected are fixed for 1hr in 4% formaldehyde/ 1x PBS, washed in 1x PBS for 2×5 minutes, cryopreserved in 30% sucrose / PBS and later frozen in OCT. Cryo-sections (6 μm) were permeabilised by incubation with 0.1% PBT for 15 minutes, followed by a heat-induced antigen retrieval step using sodium citrate buffer. Successive steps were the same as for paraffin sections (as described above).

Whole-mount staining of the cornea was performed for analysing tight junction marker ZO-1. Enucleated eyes were fixed for one hour in 4% PFA in 1xPBS. Corneal buttons are separated from the remaining parts of the eye and washed for 3×5 minutes in 1xPBS. Then permeabilised with 0.3%PBT (PBS with 0.3% triton x-100) for 1 hour, blocked with 10% BSA / 0.1% PBT for1 hour, and further antibody incubation steps were the same as for corneal sections. Then, corneal buttons are radially cut to flatten and mounted. The primary antibodies used in this study were listed in **Table S1** and secondary antibodies fused to Alexa fluorophore (Molecular Probes, 1:500). All stained sections were analysed with Zeis Imager Z2 (Zeis, Germany).

#### Phalloidin Staining

Fresh frozen samples were cut at 6μm thickness, air-dried for 30 minutes, fixed for 8 minutes at 4°C, washed in 1xPBS for 2× 5minutes, permeabilised with 0.5% PBT for 15 minutes, blocked with 10% BSA) / 0.1% PBT for 30 minutes, followed by overnight incubation with Pax6 primary antibody in 1% BSA / 0.1% PBT. Sections were washed with 0.1%PBT for 3×10 minutes, incubated with a solution containing secondary antibody (Anti-rabbit Alexa flour 488) and Alexa flour 594 Phalloidin (Invitrogen, A12381) in 1% BSA / 0.1% PBT for 1 hour and washed with 0.1% PBT, 2×10 minutes, counterstained with DAPI, washed with 1x PBS 2× 5 minutes, and mounted in Mowiol.

#### BrdU labelling

To determine proliferation in wild type and mutant mice, animals of desired age were injected intraperitoneally with BrdU (Sigma B9285, 0.1mg/ g body weight) and sacrificed by cervical dislocation after 2 hours. Enucleated eyes were fixed in 4% formaldehyde and processed to embed in paraffin wax. Paraffin sections were cut with 6 μm thickness, dewaxed and hydrated with ethanol series. The antigen retrieval step was done by heating in sodium citrate buffer (10mM sodium citrate buffer, 0.05% Tween 20, pH-6.0), followed by incubation in 2N HCl for 30 minutes and 20 minutes neutralisation using 0.1M borate buffer (pH 8.3). Subsequently, washed with PBT for 10 minutes, blocked with 10% BSA for 1 hour and incubated at 4°C overnight with rat anti-BrdU antibody (1:500, Abcam, ab6326) and rabbit anti-Pax6 in 1% BSA in PBT. Follow up steps are the same as described above for paraffin sections.

The ration of the number of proliferating cells and the number of DAPI^+^ cells for each genotype were always calculated by counting 5-6 central sections from at least three different samples. The statistical significance was determined by Student’s t-test.

## Results

### Pax6 deletion using *Aldh3-Cre* results in an abnormal cornea with reduced thickness

For conditional inactivation of Pax6 during postnatal development of CE, we used *Aldh3-Cre* transgenic mice [46] in which Cre recombinase expression is controlled by the cis-regulatory regions of the mouse *Aldh3* gene within a large BAC clone **(Fig. S1)**. To visualise Cre recombinase activity, we bred *Aldh3-Cre* transgenic mice with *Rosa26R* reporter strain [47]. Consistent with the endogenous expression of Aldh3 [48], apparent β-gal activity was found in CE from PN12-PN14, with a robust increase upon eyelid opening **(Fig. S1)**. A weak or no β-gal activity was found in limbal epithelial cells (LE) at all experimental stages **(Fig. S1)**. To inactivate Pax6 in postnatal CE, *Aldh3-cre* mice were bred with *Pax6^fl/fl^* [32]. For the sake of simplicity, we refer *Pax6^fl/fl^* as wild type and *Aldh3-Cre; Pax6^fl/fl^* as CE cKO mutant in the text.

To assess the efficiency of *Aldh3-Cre* in Pax6 inactivation, we collected eyeballs from wild type and CE cKO and analysed them by Pax6 immunostaining. A schematic representation demonstrating the Pax6 deletion pattern at different postnatal stages of corneal development is shown in **Fig. 1**. At PN12 in CE cKO mutant, reduced and patchy Pax6 expression was observed in central CE **(Fig. 1A’).** By PN21, very few or no cells express Pax6 in the central cornea **(Fig. 1B’).** At PN28, mosaic expression was observed, with no Pax6 proteins found in the peripheral cornea **(Fig. 1C’, Fig. S1)** and a patch of Pax6 positive cells at the central cornea **(Fig. 1C’, Fig. S1)**. Pax6 expression was retained in all these stages in LE and at the very end of peripheral cornea **(Fig. 1A’, B’, C’, Fig. S1)**. High Pax6 protein expression was observed in CE and LE of wild type **(Fig. 1A, B, C, Fig. S1).**

**Figure 1:**
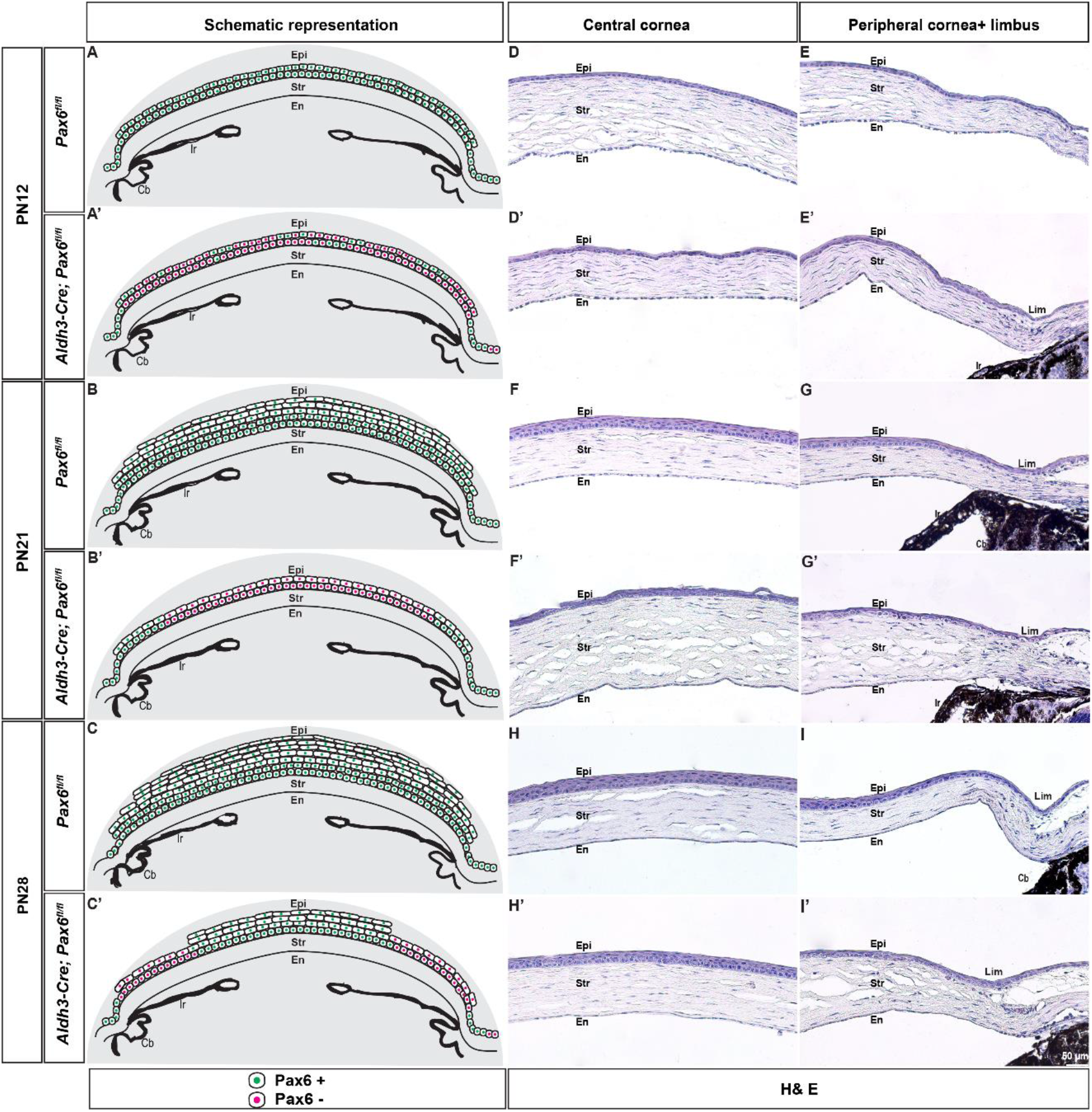
Schematic representation of *Aldh3-Cre* mediated Pax6 inactivation and morphological consequences in postnatal corneal epithelial cells. **(A-C’)** Schematic representation of Pax6 deletion pattern at indicated developmental stages. **(D-I’)** Hematoxylin and Eosin staining in frontal sections of wildtype (*Pax6^fl/fl^*) and CE cKO (*Aldh3-Cre; pax6^fl/fl^*) mutant. **(D-E’)** At PN12, no obvious difference is found in CE morphology. **(F-G’)** At PN21, CE of CE cKO mutant has less number of layers and abnormal stratification. **(H-I’)** At PN28, **(H, H’)** central cornea of CE cKO mutant is little bit thin in comparison to age-matched control, but with proper stratification as in wildtype. **(I, I’)** The peripheral cornea remains thin and with abnormal architecture. Abbreviations used in this figure and successive images: En, Corneal endothelium; Str, Corneal stroma; Epi, Corneal epithelium; Cb, ciliary body; Ir, Iris; Lim, Limbal epithelium. Scale bar: **(D-I’)**-50μm

To investigate the phenotypic consequences in response to Pax6 inactivation, we next performed histological analysis. Hematoxylin and Eosin staining revealed no significant difference between wildtype and CE cKO mutant at PN12. **(Compare Fig. 1D, E with D’, E’)**. At PN21, the wild type has 4-6 layered stratified CE with three types of cells; basal cells, wing cells and superficial cells. Basal cells are cuboidal or columnar with oval nuclei, wing cells are polygonal cells with central rounded nuclei, and superficial cells exhibit flat polygonal morphology as well as flat nuclei **(Fig. 1F, G)** [1]. At the same time point, CE cKO mutant had 2-3 layers lacking any neatly ordered architecture **(Fig. 1F’, G’)**. At PN28, the wild type CE has 5-6 layers **(Fig. 1H, I)**, whereas the central CE (which retains Pax6) of CE cKO mutant has 3-4 layers but remains morphologically similar to wild type. **(Fig. 1H’).** In contrast, the peripheral cornea (with loss of Pax6 expression) showed abnormal epithelial cells **(Fig. 1I’)**. Furthermore, histology revealed disturbed stroma with an increased number of keratocytes at the boundary between the region with and without Pax6 **(Fig. S2)**. The CE did not haven mucin containing goblet cells **(Fig. S2),** while on the contrary, goblet cells were present in central and peripheral CE in some CE cKO mutant **(Fig. S2)**. Taken together, normal corneal development is disrupted following the onset of Cre expression and depletion/reduction of Pax6 proteins marked by reduced cell layers and abnormal cell morphology.

### Pax6 loss in CE altered the differentiation status

To further characterize the abnormal corneas, we next analysed the expression of epithelial differentiation markers. Cytokeratin 12 (K12) is a bona fide marker for CE, not for basal limbal and conjunctival epithelial cells [49, 50]. In wild type, K12 is expressed in all layers of CE **(Fig. 2A, B)**. However, by PN21, K12 is lost in the entire CE of CE cKO mutants **(Fig. 2A’, B’)**. We also examined the expression pattern of Cytokeratin 14 (K14), a basal cell marker [51]. In contrast to wild type CE, where K14 is present only in the basal layer **(Fig. 2C, D),** CE cKO mutants showed strong expression in all existing layers **(Fig. 2C’, D’).** Moreover, there was no change in the expression pattern of Keratin 4 (K4) and Keratin 10 (K10), markers of the conjunctival epithelial cells [51] **(Fig. 2E-F’)**, and skin, respectively **(Fig. 2G-H’)**. Together, this data suggests that upon loss of Pax6 around the time of eyelid opening, differentiation program in CE were affected, but no changes to conjunctival or epidermal fates were found.

**Figure 2:**
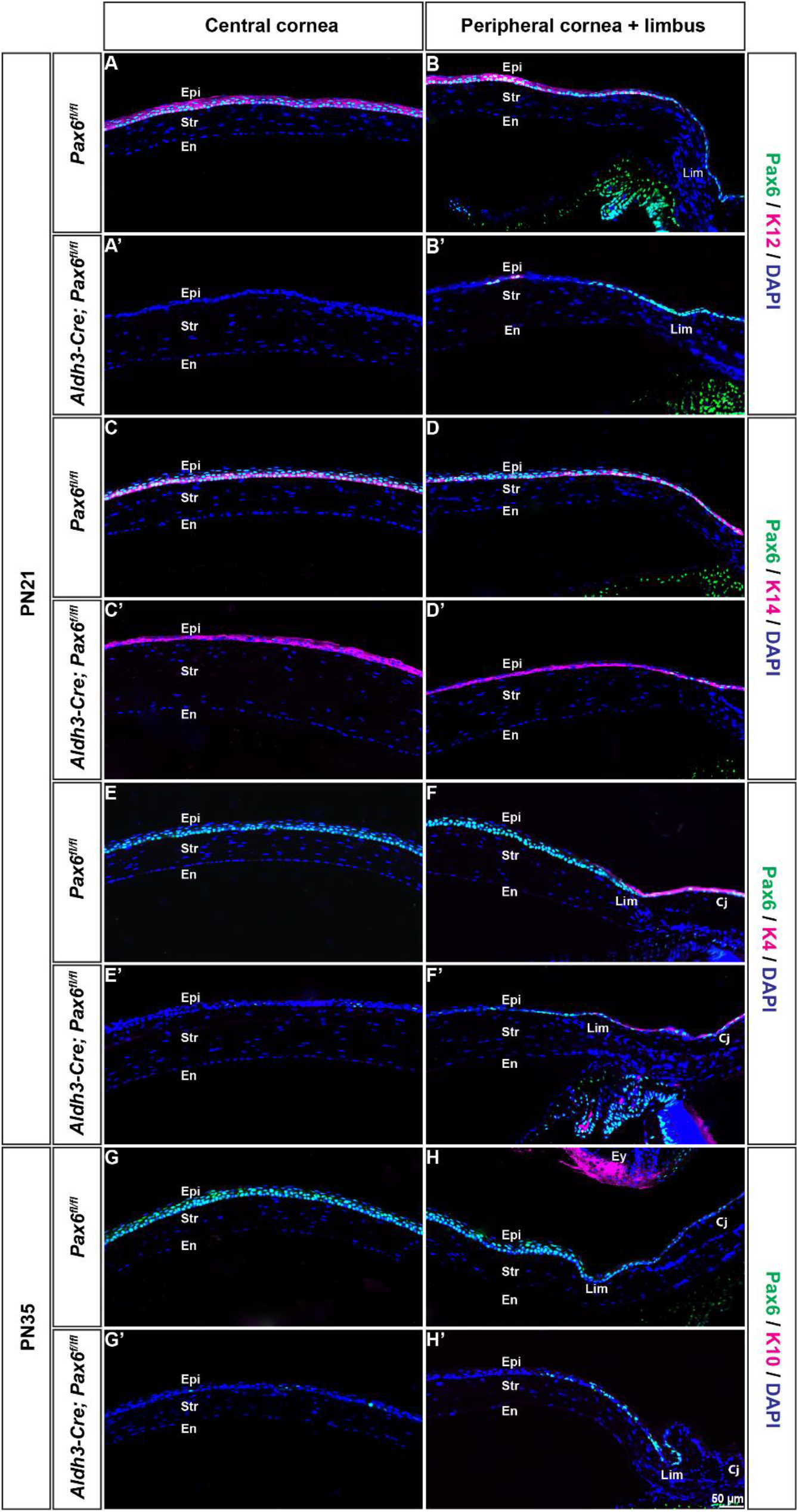
Altered expression of keratins in corneal epithelium (A-H’) Coronal sections from PN21 and PN35 of wildtype (*Pax6^fl/fl^*) and CE cKO mutant (*Aldh3-Cre; Pax6^fl/fl^*) were subjected for immunostaining with antibodies indicated. **(A, B)** Wild type CE is stained with K12, while expression of K12 is lost in **(A’, B’)** CE cKO mutant. Expression of K14 is seen only in the basal layer of **(C, D)** wildtype cornea, whereas in **(C’, D’)** CE cKO mutant K14 is detected at other layers. **(E-F)** As in control, conjunctival specific expression of K4 was not expanded to the CE in **(E’-F’)** CE cKO mutant. No immunoreactivity for K10 was found in the CE of **(G, H)** wildtype and **(G’, H’)** CE cKO mutant. Abbreviations: Cj, Conjunctival epithelium; El, Eyelid, Scale bar: **(A-H’)** - 50μm

### Pax6 loss in CE resulted in increased proliferation

Disruption of corneal epithelial differentiation and reduced thickness might be due to changes in the cell proliferation potential. To probe this directly, we investigated the proliferation status of CE at different postnatal development stages by BrdU incorporation for 2 hours. The distribution of BrdU^+^ nuclei was restricted to the basal layer of the CE **(Fig. 3B, C)**. The fraction of proliferating cells was counted by dividing the number of BrdU^+^cells by the DAPI^+^ cells in the basal layer. The number of proliferating cells was slightly increased in CE cKO mutant compared to wildtype at PN12 and PN14 (**Fig. S3**). Moreover, at PN21 and PN28, only 7.4 ± 1.2%, 5.9 ± 1.7% were BrdU^+^ in wildtype CE compared with 19.6 ± 2.1%, 20.4 ± 4.2% BrdU^+^ in CE cKO respectively **(Fig. 3A-F’, G)**. Collectively, these findings suggest that inactivation of Pax6 results in increased proliferation though this change in proliferation status does not contribute to the reduced thickness or impaired epithelial differentiation.

**Figure 3:**
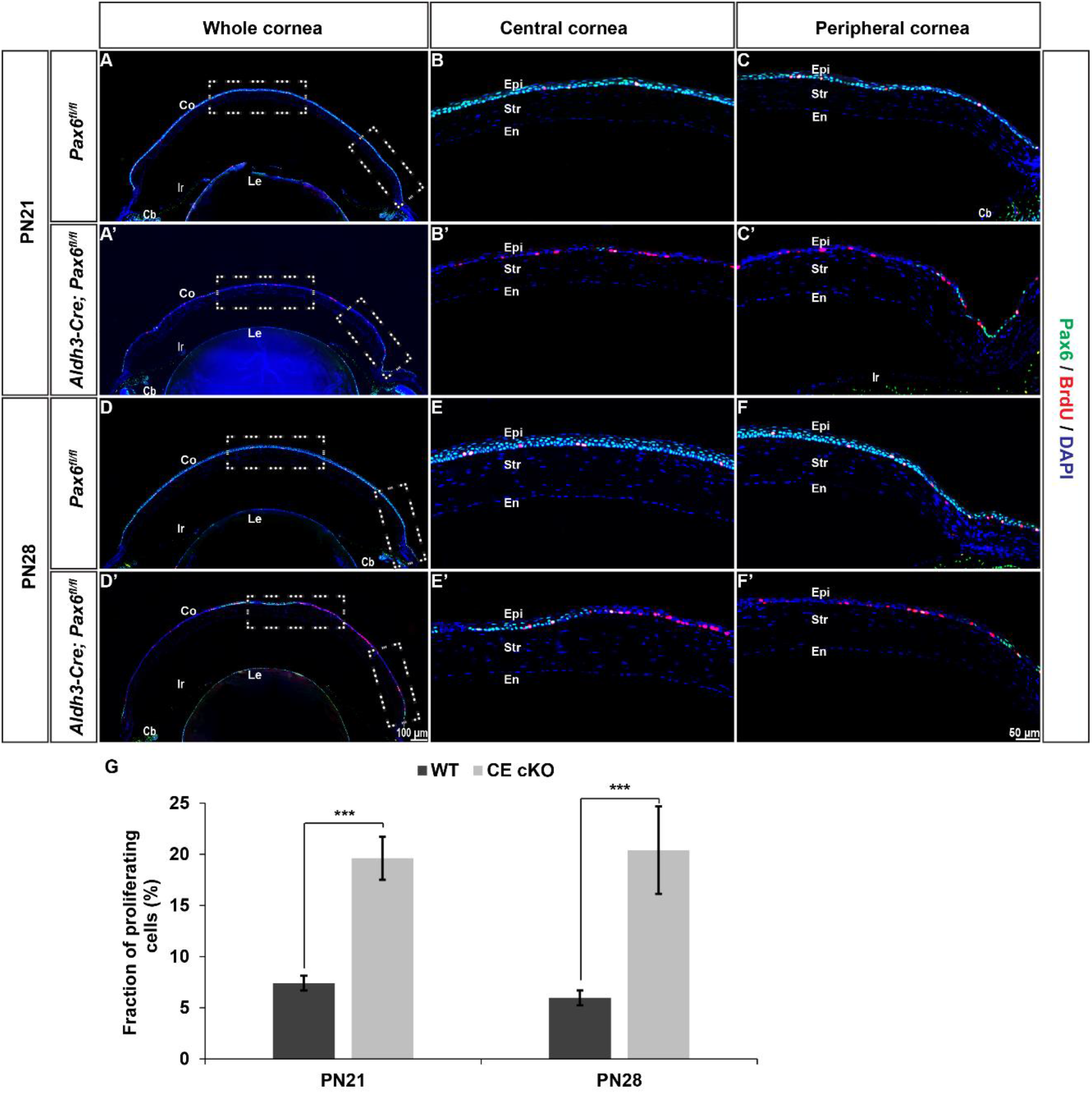
Increased cell proliferation in corneal epithelium. (A-F’) Immunostaining of Pax6 and BrdU on frontal sections of wildtype (*Pax6^fl/fl^*) and CE cKO (*Aldh3-Cre; Pax6^fl/fl^*)at PN21 and PN28. CE of **(A’-C’), (D’-F’)** Pax6 CE cKO mutant had an increased number of BrdU^+^ cells than age-matched control at (**A-C**) PN21 and (**D-F**) PN28. **(G)** The fraction of proliferating cells determined by dividing the number of BrdU^+^ cells against the total number of DAPI ^+^ cells in the basal layer. Error bars indicate standard deviation. The p-values are calculated by Students t-test. ***p < 0.001 **(A, A’, D, D’)** T^i^he dashed rectangle in the whole cornea indicates the region of interest showed in higher magnification panels. Abbreviations used; Co, Cornea; Le, Lens. Scale bar: 50μm except for **(A, A’, D, D’)** −100μm.

### Pax6 loss in CE resulted in defective cell-cell adhesion

Considering the above paradox that proliferation is increased despite the reduced thickness, we next investigated these affected cell populations at the level of cell-cell adhesion. First, we looked at the expression of E-cadherin, a key transmembrane protein of adherens junction. Immunostaining revealed abundant and proper membrane localisation of E-cadherin in wildtype CE **(Fig. 4A-B’)**. In contrast, the CE cKO mutants showed diffused expression pattern without proper membrane localisation **(Fig. 4C-D’).** Next, we analysed β-catenin, a binding partner of E-cadherin. Similar to E-cadherin, β-catenin has a plasma membrane localisation in wildtype CE **(Fig. 4I-J’),** whereas its presence sharply decreased in CE cKO mutants **(Fig. 4K-L’)**. Furthermore, we analysed the organisation of the F-actin cytoskeleton in CE using fluorescently labelled phalloidin. In contrast to the proper organisation as in wildtype **(Fig. 4E-F’)**, CE cKO mutants display more diffused expression patterns **(Fig. 4G-H’)**. Finally, we assessed the expression of tight junction protein ZO-1, which plays a vital role in the epithelial barrier function of the cornea [52, 53]. In wildtype cornea, ZO-1 has a robust continuous expression at cell-cell borders of superficial layers (**Fig. 4M, M’**), whereas CE cKO mutants have decreased expression at cell-cell borders (**Fig. 4N, N’**). These findings suggest the loss of epithelial barrier integrity upon Pax6 depletion. Our aggregate data support the model in which postnatal inactivation of Pax6 results in the loss of proper cell-cell adhesion in CE, which in turn could account for the reduced number of layers.

**Figure 4:**
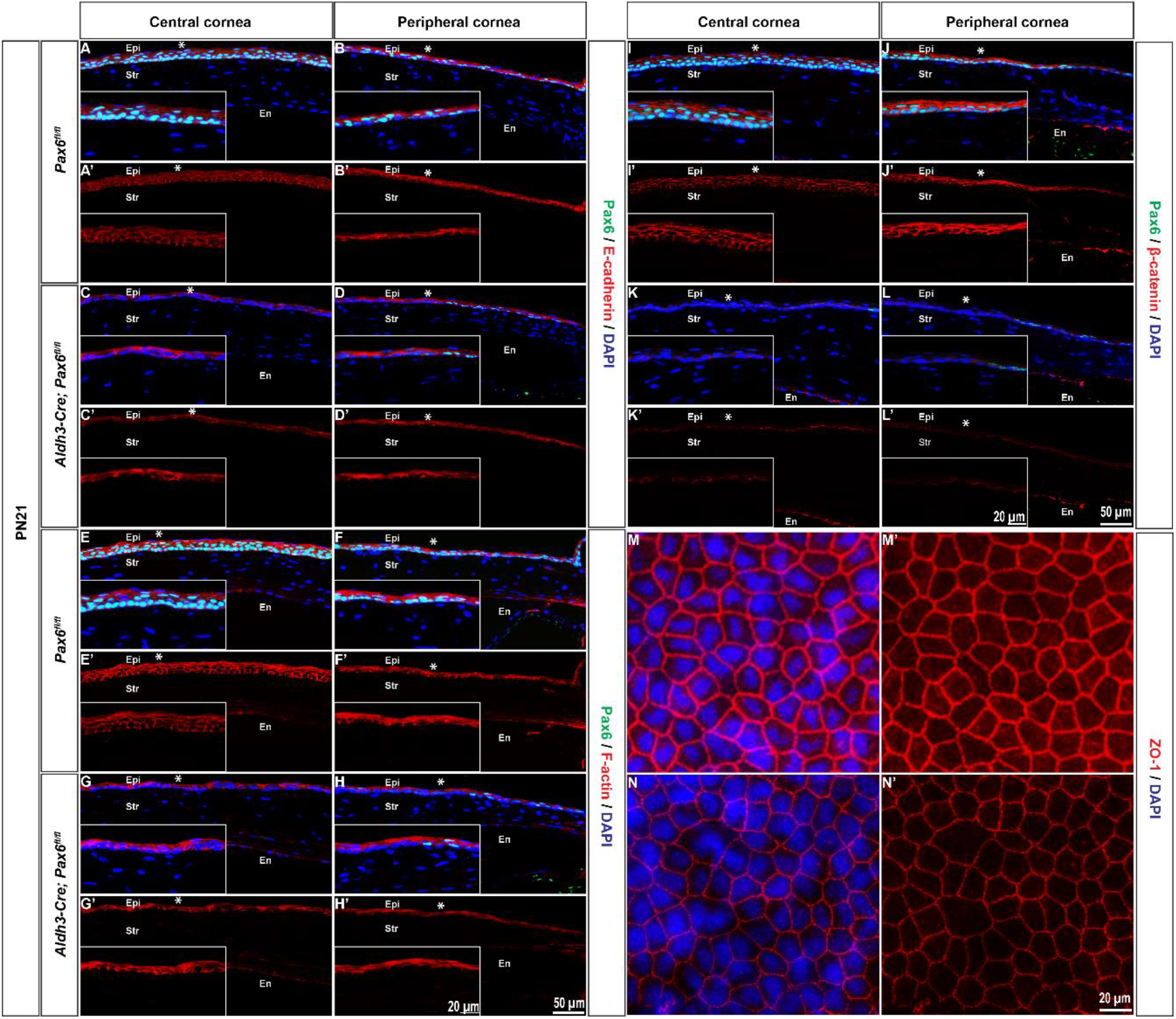
Cell adhesion and cytoskeletal defects in corneal epithelia. **(A-L’)** Coronal sections from PN21 of control (*Pax6^fl/fl^*) and CE cKO (*Aldh3-Cre; Pax6^fl/fl^*) was subjected to immunostaining with antibodies indicated. **(A-B’)** Proper membranous localisation of E-cadherin in CE cells of control eyes **(C-D’)**, while a diffused pattern was observed in CE cKO mutants. **(E-H’)** Diffused expression of F-actin is observed in CE cKO mutant in contrast to its proper organisation in control. **(I-J’)** Control CE cells have membrane localisation of β-catenin, whereas (**K-L’**) CE cKO mutant has an overall downregulated expression in CE cells. **(M-N’)** Whole-mount immunostaining of ZO-1 in superficial layers of CE at PN21. Inserts are higher magnification images of the region of interest indicated by asterisks **(*)**. Scale bar: **(A-L’)**-50μm and inserts - 20μm, **(M-N’)** −20μm

### Pax6 loss in the OSE using *K14-Cre* resulted in thickened epithelia

Next, for conditional inactivation of Pax6 in OSE from embryonic stages, we used *K14-Cre [45]*, where Cre recombinase activity is controlled by the K14 promoter. To visualise the Cre activity, we crossed the *K14-Cre* transgenic mice with *Rosa26R* reporter strain [47]. X-gal staining revealed weak reporter activity in CE, while modest Cre activity is observed in the presumptive conjunctival epithelium during embryogenesis (**Fig. S4**)

To generate conditional knockout mutant *K14-Cre; Pax6^fl/fl^*, we crossed *K14-Cre* transgenic mice with *Pax6^fl/fl^*. Hereafter, we refer *K14-Cre; Pax6^fl/fl^* as OSE cKO mutant and *Pax6^fl/fl^* as wildtype in the text. To investigate the efficiency of the *K14-Cre* recombinase system in inactivating the Pax6 gene, we performed Pax6 immunohistochemistry. Consistent with Cre expression, patchy Pax6 protein expression was observed in presumptive CE of OSE cKO at E15.5 and E18.5 **(Fig. S4)**. In contrast from E15.5 CE, Pax6 expression was decreased in presumptive conjunctival epithelial cells of OSE cKO mutant **(Fig. S4)**. At PN2, an asymmetrical deletion pattern was observed in most of the OSE cKO mutants; one periphery has loss of Pax6^+^ cells, while on the contrary, the other periphery and conjunctiva retains Pax6 expression **(Fig. 5A-C’)**. By PN6, very few or no Pax6^+^ cells were found in the entire OSE domain **(Fig. 5D-F’).**

**Figure 5:**
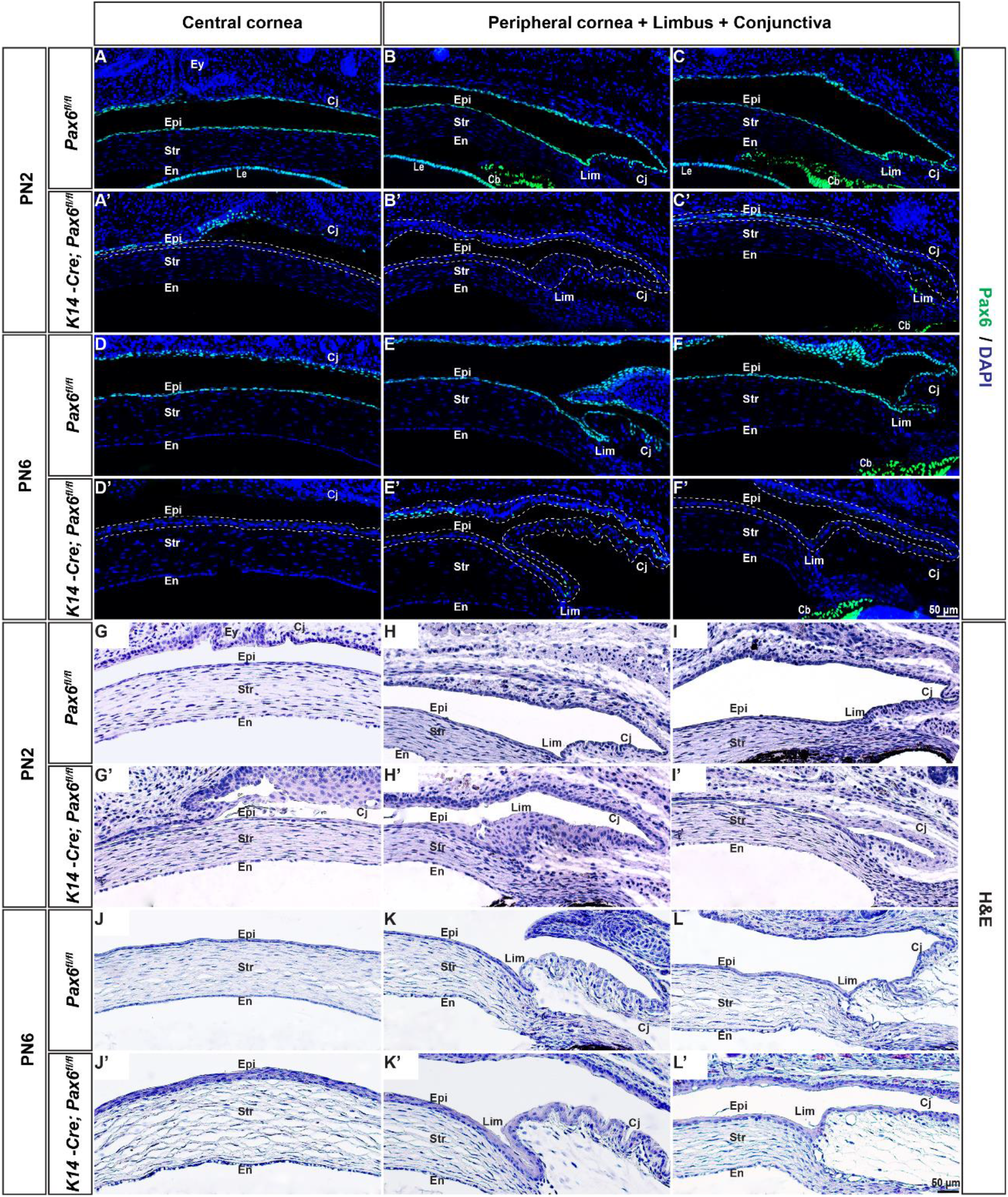
The *K14-Cre* mediated deletion of Pax6 and histological changes in the ocular surface epithelium. **(A-L’)** Consecutive frontal sections of control (*Pax6^fl/fl^*) and OSE cKO mutants (*K14-Cre; Pax6^fl/fl^*) were stained with Pax6 and H&E at indicated postnatal stages. **(A-C’)** At PN2, Pax6 expression is retained in the one peripheral part of the cornea while it is deleted in other periphery and central epithelial cells. **(D-F’)** At PN6, Pax6 expression is absent in the central, limbal and conjunctival epithelial cells of OSE cKO mutants. **(G-I’)** At PN2, OSE cKO mutants have thicker LE compared to control. **(J-L’)** At PN6, OSE cKO mutant exhibit moderately thickened corneal, limbal and conjunctival epithelium compared to control epithelium. The dashed line indicates the region of Pax6 deletion. Scale bar: **(A-L’)**-50μm

To further characterise the phenotypic changes, we next examined morphological and histological changes in OSE cKO mutants and age-matched controls. Hematoxylin and eosin staining at PN2 revealed thickened limbal and conjunctival epithelium in OSE cKO mutant **(Fig. 5H’)** in contrast to 1-2 layered wildtype epithelia **(Fig. 5H, I)**. Simultaneously, part of the cornea and limbus which retains Pax6 remained thin **(Fig. 5I’)**. Besides that, no significant change is found in the central CE **(Fig. 5G, G’)**. At PN6, in comparison to wildtype tissue morphology, the corneal, limbal and conjunctival epithelia of the OSE cKO mutants are moderately thickened **(Fig. 5J-L’)**. By PN18, palpebral spaces in OSE cKO mutants are small with some of them have eyelashes touching the CE **(Fig. S5)**. By PN23, eyes of OSE cKO mutants appear opaque with 20% penetrance **(Fig.S5)**. Histological analysis of opaque eyes at PN23 showed keratinisation and extensive thickening of central CE of OSE cKO compared to age-matched control animals **(Fig. S5)**. Collectively, these data show that Pax6 inactivation in OSE during early postnatal stages impairs ocular surface homeostasis.

### Pax6 loss in OSE resulted in CE adopting keratin expression in conjunctival epithelia

Our present histological analysis indicates a cell fate switch in the ocular surface. Our hypothesis predicts that loss of Pax6 affects the cell fate decisions in the OSE compartment and is thus involved in major corneal postnatal developmental processes. To test this hypothesis, we examined the expression of epithelial markers at early postnatal stages (PN1-PN12) prior to eye-opening. In contrast to wildtype eyes, K12 expression is lost in OSE cKO mutants **(Fig. 6A-C’)** while expression of K14 expands to all cell layers in the cornea by PN6 **(Fig. 6D-F’)**. These findings demonstrate that corneal epithelial differentiation is impaired upon Pax6 inactivation. In control conjunctiva, K14 is expressed in basal and suprabasal cells **(Fig. 6D-F)**, OSE cKO mutants maintain the same expression pattern even though they have comparatively thicker epithelium **(Fig. 6D’-F’)**. In most cases, CE of OSE cKO mutants is devoid of K10 during early postnatal stages **(Fig. 6G-I’)**. A rare exception, with patches of K10^+^ cells, was observed in samples with comparatively no conjunctival sac and had more or less direct contact of CE with thickened conjunctiva (**Fig. S6**). Keratinisation in these samples could be a response to irritations caused by the closeness of thickened conjunctival epithelia and CE. Therefore, we further analysed expression of K4, which is localised to the conjunctival epithelium in wildtype mice. K4 expression expanded into both the CE and LE in OSE cKO mutants **(Fig. 6J-L’)**. Based on this data, we infer that the loss of Pax6 in OSE during early postnatal stages result in conjunctivalisation of CE.

**Figure 6:**
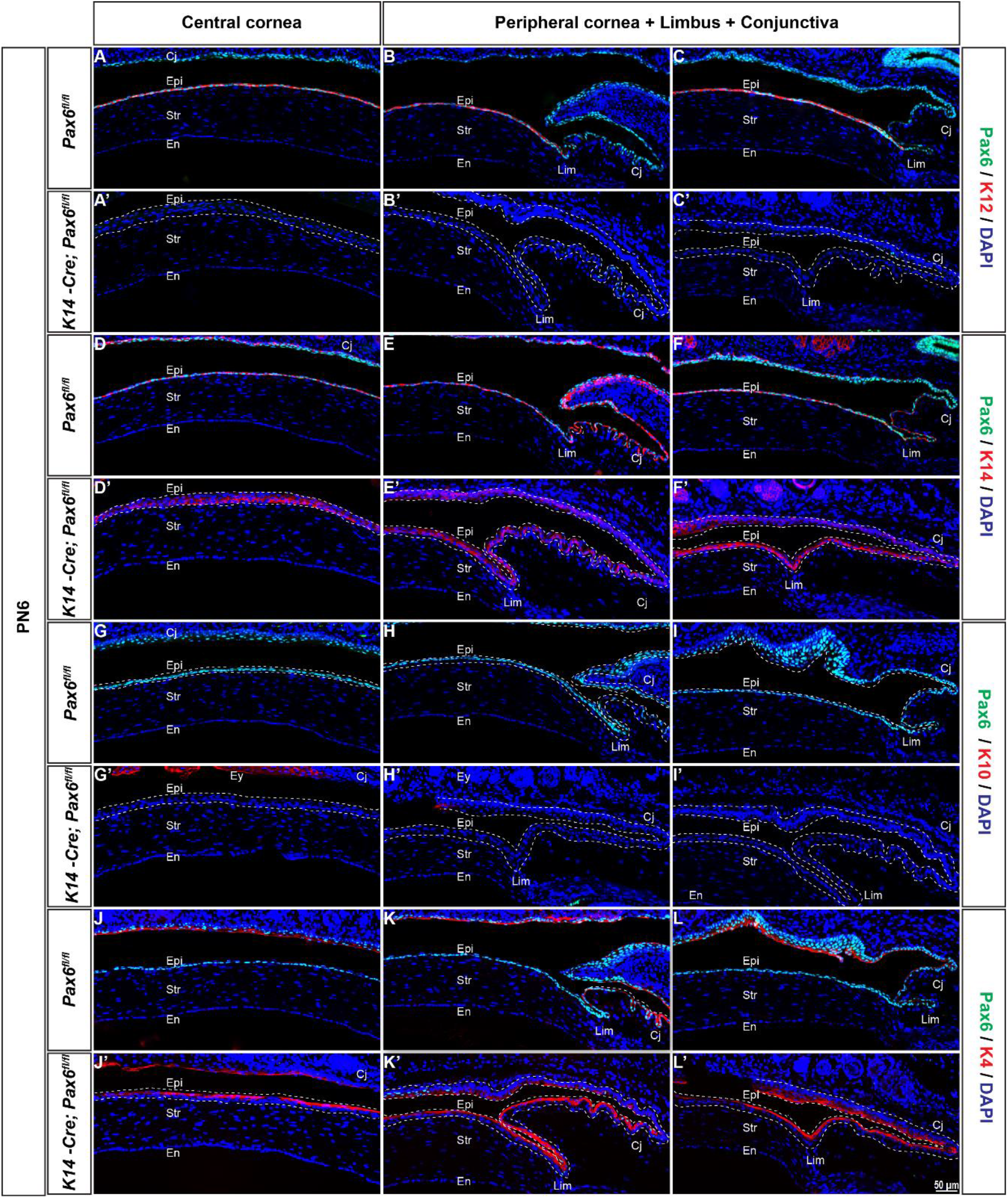
Keratin expression in the CE adapts pattern in conjunctival epithelium until eye opening. **(A-L’)** Coronal sections of control (*Pax6^fl/fl^*) and OSE cKO mutants (*K14-Cre; Pax6^fl/fl^*) were stained with indicated antibodies**. (A-C’)** K12 expression is lost in CE cells of OSE cKO mutants. **(D-F’)** K14 expression is expanded apically to all cell layers in OSE cKO mutants, while it is restricted to the basal cell layer in control CE. **(G-I’)** No immunostaining of K10 is observed either in the CE of OSE cKO mutant or control. **(J-L’)** Expansion of K4 expression to the CE in the OSE cKO mutant, whereas it is confined to the conjunctival epithelium in control. Dashed lines indicate OSE with Pax6 loss in OSE cKO mutants. Scale bar: **(A-L’)** −50μm

To complete these analyses, we examined expression of three epithelial markers, including K12, K14 and K10, at PN23. Expression of K12 is absent in thickened epithelia of OSE cKO mutants, while the expression of K14 is expanded to apical cell layers in OSE cKO mutants in contrast to its basal layer expression in control eyes **(Fig. S6)**. Furthermore, expression of skin-specific K10 was observed in CE with Pax6 loss **(Fig. S6)**. Even though this data indicates the developmental fate change of CE to the epidermis at PN23, we interpret this fate change as a response to irritations happening to the corneal surface, including small palpebral space, infection, and contacts with eyelashes.

## Discussion and conclusions

This is the first study to define roles of Pax6 in postnatal mouse corneal development and to integrate these results with previously established critical functions of this transcription factor in early corneal development. These findings are both important for our understanding of intricate complexity of corneal development and maintenance and pathology of cornea in human aniridia patients [36, 37]. To accomplish these goals, we employed genetic conditional loss-of function approach using two stage- and cell-specific *Krt14*- and *Aldh3-cre* lines. Previous studies were limited to analysis of various Pax6 haploinsufficiency models [24, 54, 55], however, in both our *Krt14* and *Aldh3-cre* models, corneal development proceeds normally till PN1 and PN12-14, respectively. Our data show that Pax6 is important for postnatal development of the cornea and the maintenance of the limbal epithelial identity.

### Role of Pax6 in CE during postnatal development

Selective inactivation of Pax6 in CE concomitant with eye-opening, leads to the abnormal cornea marked by only 3-4 cell epithelial layers concomitant with altered cell-cell adhesions in the CE of CE cKO compared to control littermates. Adherens junctions are multi-protein complexes essential during development and tissue morphogenesis [56, 57]. E-cadherin is a transmembrane protein, a core component of adherens junctions [58, 59]. The cytoplasmic tail of cadherin interacts with catenin proteins which further links cadherins to the actin cytoskeleton [60–62]. The CE has E-cadherin expression in all layers and is known to be vital for the maintenance of its structural integrity [59, 63]. The initial analysis of cornea using chimeric AP-2α wild type and null cells showed downregulation of E-cadherin in mutated cells [64]. In the follow up studies, E-cadherin expression was decreased in the lens and CE of AP-2α conditional knockout mutants [65]. Our study found that abundant membrane localisation of E-cadherin in CE is lost in CE cKO mutants. A similar diffused pattern of expression is observed for F-actin and β-catenin. This suggests that adherens junctions are aberrant in CE upon Pax6 loss, accounting for the reduced number of cell layers.

Previous studies have shown that retinoic acid-responsive transcription factor AP-2α together with Pax6 control gelatinase B (Mmp9) promoter activity in response to wound healing [66]. AP-2α has an overlapping expression with Pax6 in both the embryonic and adult eye [67, 68]. Postnatally, like Pax6, AP-2α was observed mainly in the basally located CE [65, 67]. *In vitro* studies proposed AP-2α mediated E-cadherin promoter activity [69, 70]. Corneas of AP-2α conditional knockout mutants exhibit similar phenotype to Pax6 deficient ones, including adhesion defective thin epithelium [24, 65]. Taken together, these findings raise the possibility of co-operative regulation of E-cadherin expression by Pax6 and AP-2α. Further studies are required to determine whether AP-2α regulates E-cadherin through its co-operative interaction with Pax6.

Interestingly, Pax6 heterozygous corneas showed no change in E-cadherin expression [24, 68]. It is likely that one copy of *Pax6* is sufficient to maintain the promoter activity of E-cadherin, either in co-operation with AP-2α [68]. This is further supported by the notion that no morphological changes were found upon conditional deletion of one copy of Pax6 in postnatal CE cells using tamoxifen-inducible Cre model [71]. However, these models delete floxed Pax6 allele using the ubiquitous CAG promoter and cell-specific Krt19 at E9.5 and 12 weeks postnatally, respectively [71].

Our CE Pax6 cKO data show for the first time that the tight junction protein ZO-1 between the superficial epithelial cells is downregulated in any Pax6 gene loss-of-function murine model consistent with increased fluorescein uptake in corneas of heterozygous Pax6 mutant mice [24], which indicates defective tight junctions upon Pax6 loss. This could also explain increased fluorescein uptake in heterozygous Pax6 mutants. The formation of adherens and tight junctions is interdependent, and loss of adherens junction components could lead to cells that do not develop tight junctions [72–75]. It is plausible that a decrease in adherens junction components could affect other cell-cell junctions and, thus, total cell-cell adhesion.

Keratin 12 (KRT12), a major intermediate filament with corneal specific expression [76], has been controlled by PAX6 together with reprogramming factors KLF4 and OCT4 [77, 78]. Consistent with this, our data indicated the loss of Krt12 in CE cKO mutants. Homozygous knockout models in mice [79] and mutations in the human KRT12 gene [80] are characterized by corneal fragility. We thus propose that loss of Krt12 in the CE cKO mutant could further deteriorate intercellular adhesion. As discussed above, expression of Krt12 indicates cornea specific differentiation, loss of Krt12 in CE cKO mutant corneas suggest impaired differentiation program. This observation is further supported by a robust and multilayered expression of basal cell-specific Krt14 in Pax6 deficient CE.

To further evaluate the possibility of transdifferentiation of CE to the skin-like cells or the migration of conjunctival cells, we examined Krt10 and Krt4 expression, respectively. Although earlier studies reported Krt4^+^ cells in the CE of heterozygous Pax6 mutants [24], we do not observe any expansion of Krt4 or Krt10 to CE. At the same time, goblet cells are observed in peripheral and Pax6 retaining regions (Krt12^+^) of the central cornea in CE cKO mutants at PN28. The presence of goblet cells in CE is considered a major characteristic of limbal stem cell deficiency [81, 82]. The stem cells in the limbus are unable to maintain epithelial homeostasis, and as a result, cells migrate from the conjunctiva into cornea [81, 82]. Goblet cells are observed before the time point at which limbal stem cells are assumed to be active [11]. Hence the ectopic goblet cells cannot be explained as a direct consequence of limbal stem cell deficiency/limbal barrier breach. Incongruent with this, a recent study reported the presence of compound niche at the limbal-corneal border, which contains a heterogeneous population of cells including slow-cycling corneal progenitor cells, proliferating cells, non-proliferating cells and post-mitotic Krt8^+^/Krt12^+/^Muc5ac^+^ goblet cells. The authors hypothesized that, in the case of wounds in the CE that occur near the limbus, goblet cells increase in number, and they arise from cells in the compound niche, not from bulbar conjunctiva [83]. So, we assume that ectopic goblet cells in the CE of CE cKO mutants could be raised from progenitors in the compound niche in response to severe wounding/erosions due to adhesion defects.

Increased proliferation in the CE cKO corneas is consistent with the previous findings from Pax6 heterozygous corneas [24, 26]. We propose that this could be for cell loss compensation due to adverse adhesion defects [84]. Alternatively, in ocular lens, Pax6 was found to control cell proliferation how and via (EDIT) targets [32, 85].

Robust expression of Aldh3/Aldh3a1 proteins in corneal epithelium is directly regulated Pax6, Oct1/Pou2f1 and p300 [48]). However, somatic depletion of Aldh3 proteins generates structurally normal corneas [86], and, thus, we did not investigate further reduction of Aldh3 proteins in our system.

In summary, our data have revealed that *Aldh3-Cre* mediated conditional deletion of Pax6, Pax6 is required for proper maintenance of strong intercellular adhesion and the differentiation of CE during postnatal development (**Fig. 7A**).

**Figure 7:**
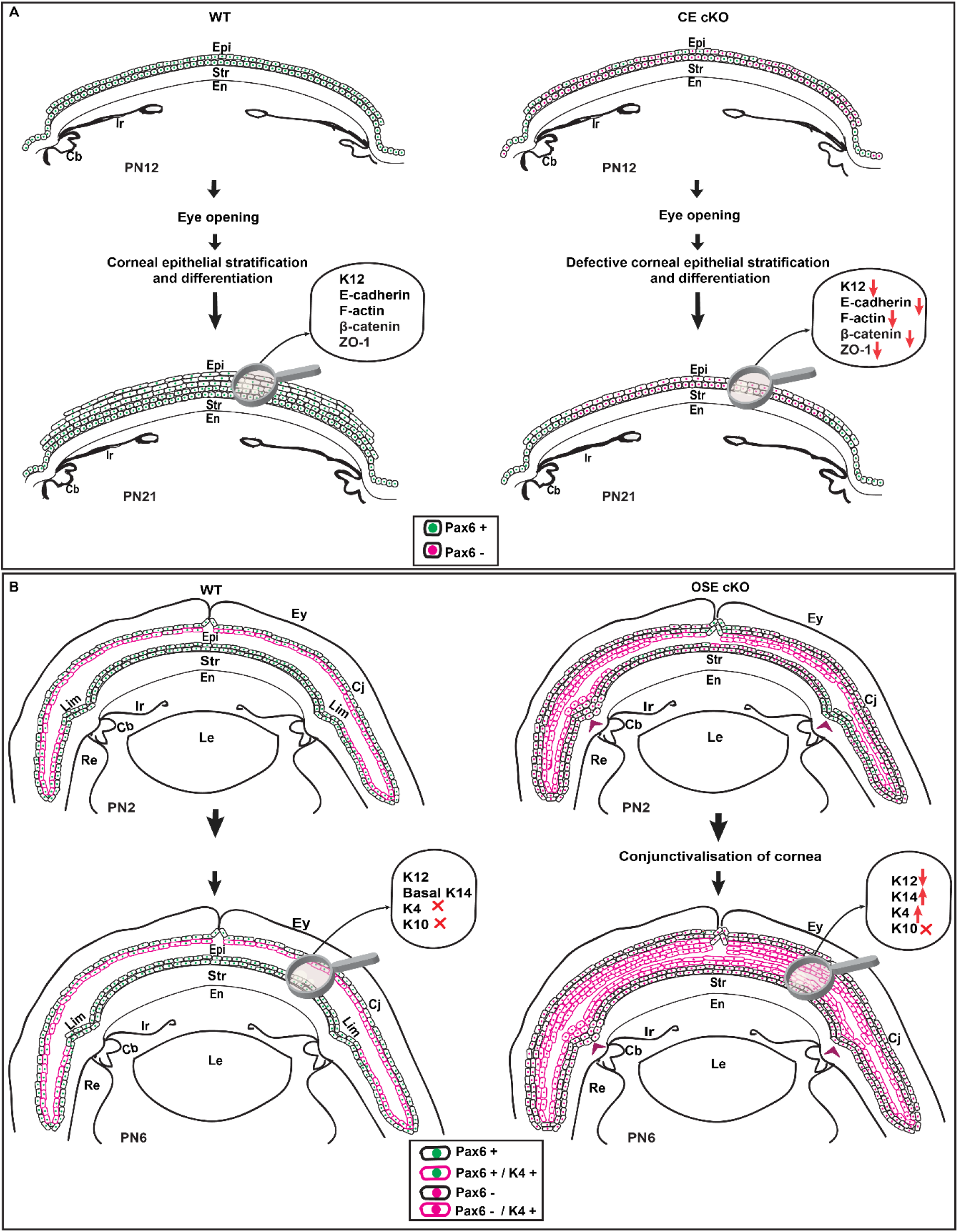
Schematic summary of the plausible role of Pax6 in developing cornea. **(A-B)** Schematic summary of the plausible function of Pax6 during mouse corneal development. (**A**) Deletion of Pax6 during postnatal corneal development in CE results in the thinner cornea with adhesion defects. **(B)** Deletion of Pax6 in OSE during early postnatal stages results in conjunctivalisation of CE cells. Arrowheads in (**B**) indicates the position of the limbus in OSE cKO mutants

### Role of Pax6 in the early postnatal OSE

To unravel the autonomous function of Pax6 in OSE during embryogenesis, we *used K14-Cre,* which is controlled by promoter-specific for basal keratinocytes. Targeted deletion of Pax6 using *K14-Cre* in OSE results in the migration of K4^+^ conjunctival cells into the cornea by PN6. Conditions in which limbus is partially or entirely deleted or lost due to chemical burns, varying degree of limbal stem cell deficiency (LSCD) arise, which results in conjunctivalisation of the cornea [50, 87–92]. Whereas in OSE cKO mutants, migration of conjunctival cells to cornea occurs prior to limbal stem cell activation [11], it is unlikely to have LSCD in our model. Here, we interpret our phenotype as a loss of limbal epithelial identity, which results in the overgrowth of conjunctival epithelial cells to the cornea.

In the anterior segment of the eye, lens and epithelial components of the eyelid, cornea, limbus, conjunctiva, lacrimal gland, and harderian gland are specified from OSE [93, 94]. Spatiotemporal control of Wnt/β-catenin signaling is important for determining these specific fates within OSE. For example, activation of Wnt/β-catenin signaling in the early OSE prevents lens induction, whereas its ablation results in ectopic lens in peripheral SE [95–97]. In addition to this, aberrant expression of β-catenin in presumptive corneal and conjunctival ectoderm disrupts its proper morphogenesis and results in ocular surface squamous metaplasia. [98, 99] Receptors and ligands of Wnt/β-catenin signaling are consistently expressed throughout the OSE [100, 101]. However, the active Wnt signaling is detected in early conjunctival epithelium and underlying mesenchyme of cornea, limbus and conjunctiva, not in presumptive corneal and limbal ectoderm [100, 102, 103]. This regional specificity could be necessary for the barrier function of limbal epithelium, and this may be maintained by the active repression of this pathway at various levels. It is further supported by the wide expression of Wnt inhibitors DKKs and Sfrp1 in the cornea during embryonic and postnatal development [100, 101, 104]. The effect of aberrant Wnt signaling depends on the stage and region of activation; CE could undergo fate switch to the epidermis [99, 104–107] dedifferentiate to more progenitor-like cells [99, 105] proliferate in an unregulated fashion [98, 99] or could inhibit stratification of CE cells [103, 108].

Pax6 directly regulates Wnt inhibitors like Sfrp1, Sfrp2 and Dkk1in the lens and central nervous system [109–111]. Moreover, elevated mRNA levels of Wnt inhibitors Wif1 and Sfpr2 were found in the cornea with Pax6 overexpression [112]. Loss of OSE identity and bias towards conjunctival fate was reported for *Dkk2^-/-^* mutants similar to our model. Taken together, we assume that Pax6 regulates the expression of Wnt inhibitor Dkk2 in the cornea. Further research needs to be carried out to see whether Dkk2 is regulated by Pax6 or not.

Finally, as we found keratinization of the cornea by PN23 in OSE cKO mutants, CE of *Dkk2^-/-^* mutants also exhibited keratinization and opaque cornea postnatally [102]. In *Dkk2^-/-^* mice, keratinization is assumed as the consequence of irritations caused by ectopic eyelashes and infections raised due to eyelid defects [102]. In our study, during early postnatal stages, ectopic K10^+^ cells are observed very rarely in OSE, and they exhibit a very narrow conjunctival sac, which could cause rubbing of CE with conjunctival epithelia. We believe that complete fate switch to the epidermis seen at PN23 in OSE cKO mutants either occurred progressively during development as a disease response to constant contact of CE with conjunctival epithelium or by chronic irritations by eyelashes after eye-opening (**Fig. S5**).

In conclusion, conditional inactivation of Pax6 in OSE using *K14-Cre* revealed the role of Pax6 in maintaining the epithelial identity of limbal cells. We observed loss of corneal epithelial identity and migration of conjunctival cells to cornea upon Pax6 loss in OSE (**Fig. 7B**).

## Acknowledgements

We thank A. Cvekl for critical comments on the manuscript. This work was supported by the grant of Czech Science Foundation (21–27364S). Mouse costs were partially covered by the Czech Center for Phenogenomics infrastructure supported by LM2018126, OP VaVpI CZ.1.05/2.1.00/19.0395, and CZ.1.05/1.1.00/02.0109.

## Competing interest

The authors declare no competing interests

## Notes

### Competing Interest Statement

The authors have declared no competing interest.

