## Supplementary file for "Multiple roles of Pax6 in corneal limbal epithelial cells and maturing epithelial cell adhesion"

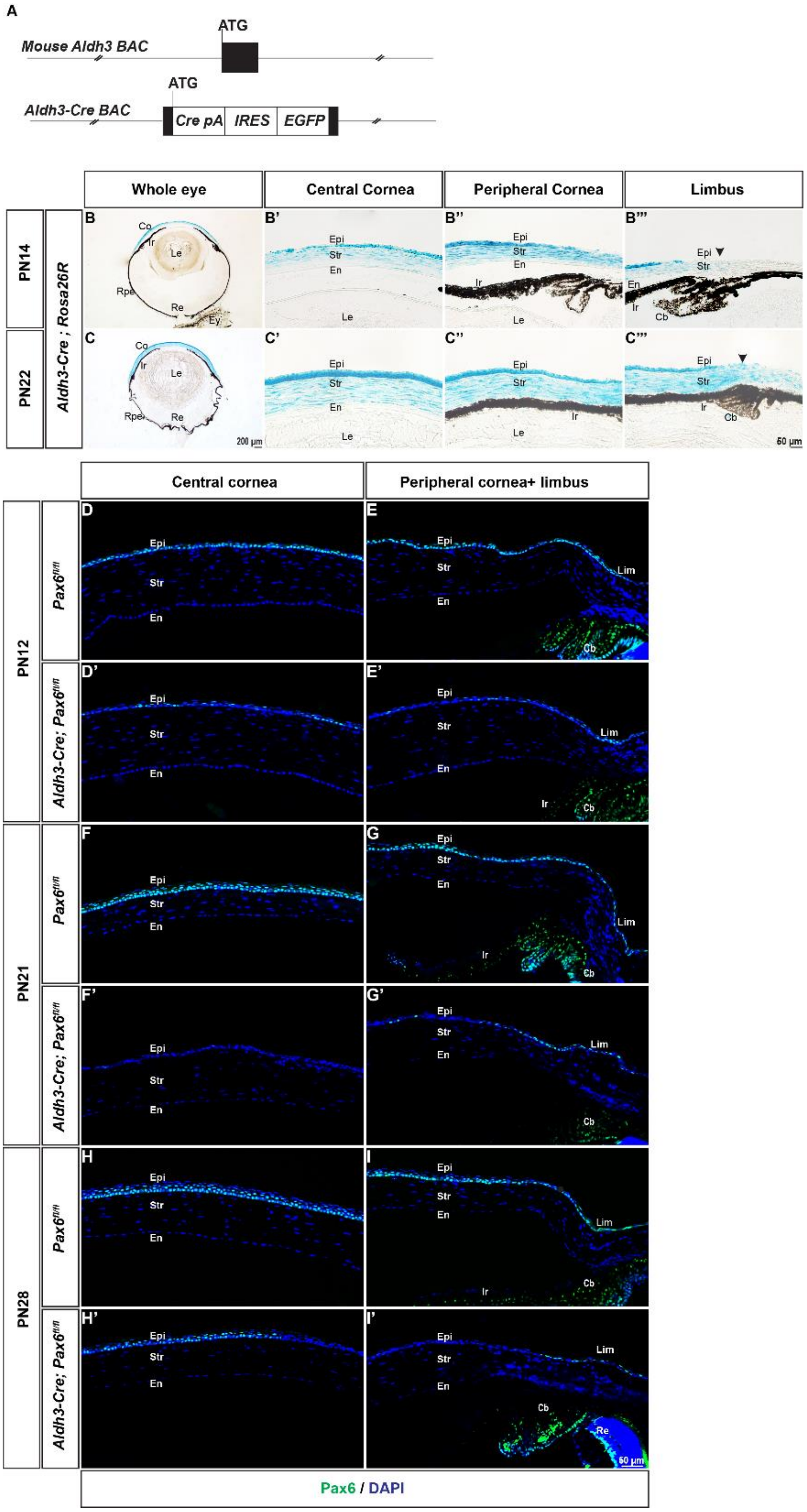


**Supplementary Figure 1: Characterisation of Cre recombinase activity of *Aldh3-Cre* transgenic mice and Pax6 deletion pattern in the cornea CE cKO mutants**

**(A)** To generate *Aldh3-Cre,* BAC containing regulatory sequences of the Aldh3 gene was modified by BAC recombineering. A cassette containing *Cre* recombinase (Cre-pA) and *EGFP* linked by Internal ribosomal entry point (*IRES*) sequence was integrated into the first translational start site (ATG) of the *Aldh3* gene. **(B-C”’)**The *Aldh3-Cre* activity was visualised using *Rosa26R* reporter strain. The β- Gal activity in frontal sections of the eye at **(B-B’’’)** PN14 and **(C-C”’)** PN22. **(D-I’)**Coronal sections of wildtype ($Pax6^{fl/fl}$) and CE cKO (*Aldh3-Cre;* $Pax6^{fl/fl}$) stained with Pax6 antibody at the indicated stages. Decreased levels of Pax6 in CE at PN12 **(D’, E’)** in CE cKO mutant comparison to **(D, E)** age-matched control**.** **(F-G’)** At PN21, very few or no active $Pax6^+$ cells are found in the CE cKO. At PN28, patchy expression of Pax6 is detected in **(H, H’)** central cornea of CE cKO, whereas deletion is observed in **(I, I’)** peripheral cornea. **(E', G', I')** At all stages, Pax6 is retained in LE cells. **(B”’, C’’’)** Black arrowheads indicate the limbus, with very low or no Cre activity. Abbreviations used: Co, Cornea; Le, Lens: Re, Retina; Rpe, Retinal pigment epithelium. Scale bar: **(B, C)** - 200µm **(B’-I’)** - 50µm.

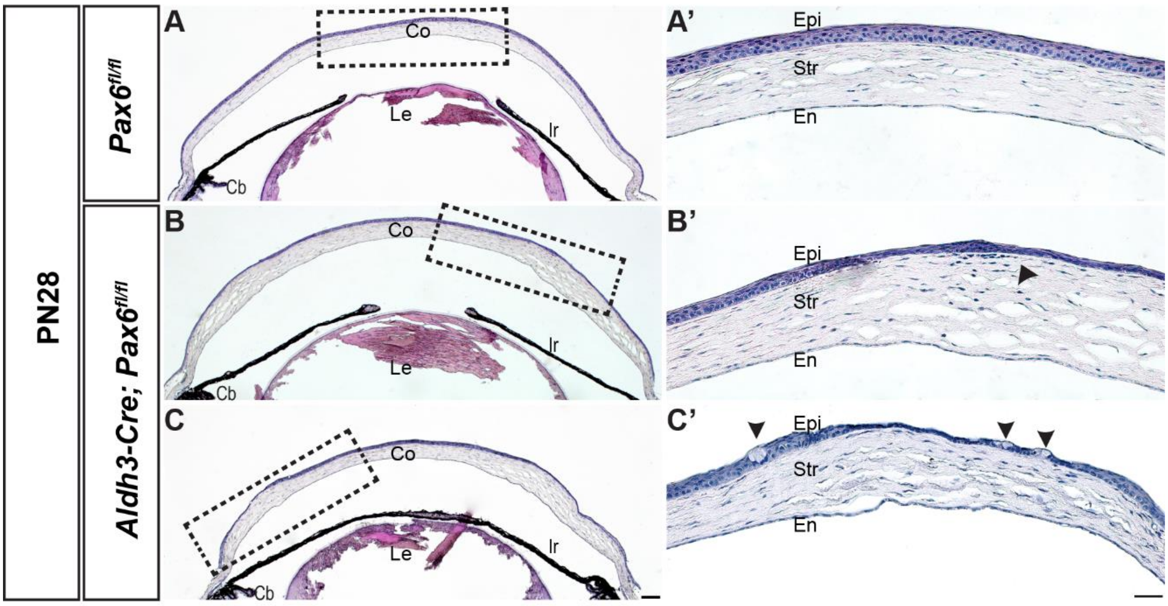


**Supplementary Figure 2: Morphological changes in the cornea at PN28**

**(A-C’)** Hematoxylin and Eosin staining of eyes of control ($Pax6^{fl/fl}$) and CE cKO mutant (*Aldh3-Cre;* $Pax6^{fl/fl}$) at PN28. **(A-A’)** In contrast to control mice, **(B-B’)** CE cKO mutant has disturbed stroma with an increased number of keratocytes at regions with loss of Pax6. **(C-C’)** Goblet cells are absent in control CE cells, while the CE cKO mutant had ectopic goblet cells in CE cells. **(A, B, C)** Dashed rectangle in the whole cornea indicates the position of the region of interest shown in higher magnification panels. Black arrowheads in **B’** and **C’** indicates keratocytes in the stroma and goblet cells, respectively. Scale bar: **(A, B, C)** - 100µm, **(A’, B’, C’)** - 50µm.

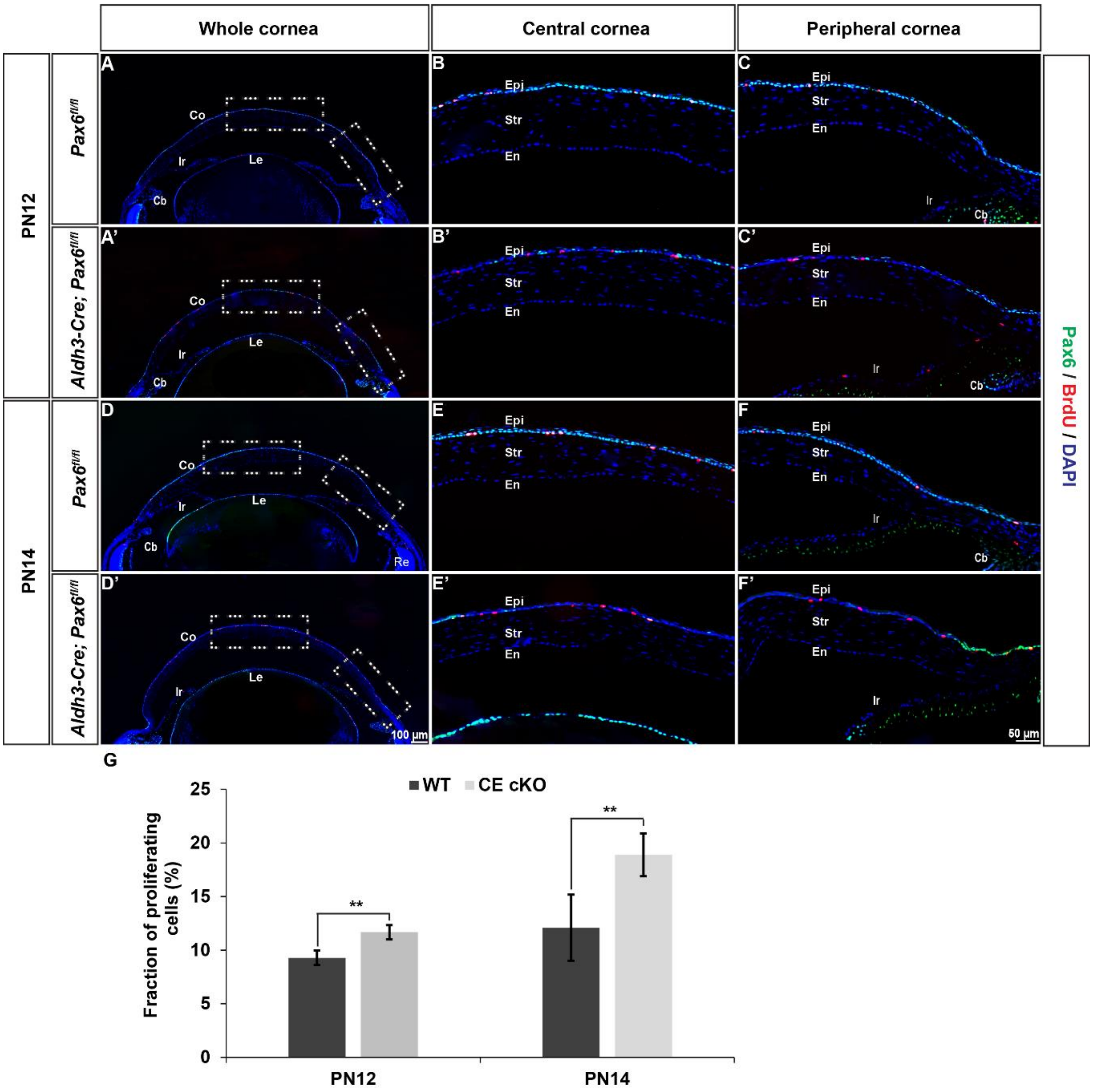


**Supplementary Figure 3: Proliferation in CE cKO mutant corneas**

**(A-F')** Coronal sections of control (*Pax6$^{fl/fl}$*) and CE cKO mutants (*Aldh3-Cre; Pax6$^{fl/fl}$*) were stained with Pax6 and BrdU antibodies at PN12 and PN14. CE of CE cKO mutants at **(A-C')** PN12 and **(D-F')** PN14 has a slight increase in the number of BrdU$^{+}$ cells in comparison to control corneas. **(G)** Quantification of S-phase cells determined by the proportion of BrdU$^{+}$ cells against the total number of DAPI$^{+}$ cells in the basal layer. Error bars indicate standard deviation. The p-values are calculated by Students t-test. ** $p < 0.01$ Scale bar: (**A-F'**) -50µm

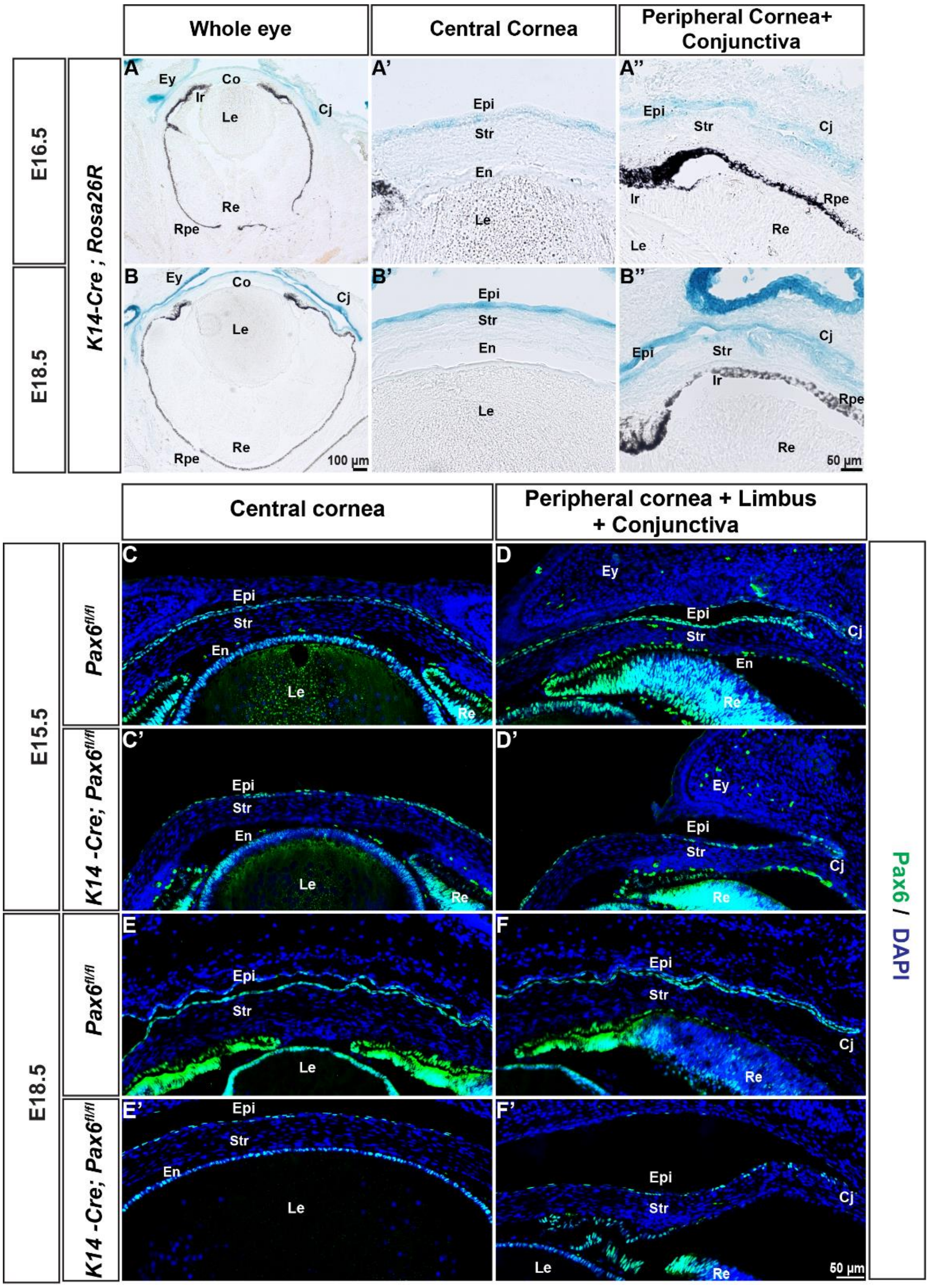


**Supplementary Figure 4: Characterisation of Cre recombinase activity in *K14-Cre* transgenic mice and Pax6 deletion pattern in the OSE of OSE cKO mutants**

**(A-B'')** The *K14 -Cre* activity was visualised by *Rosa26R* reporter strain. X-gal staining in coronal sections at (**A-A''**) E16.5, (**B-B"**) E18.5. **(C-F')** Pax6 immunostaining in the eyes of OSE cKO mutants (*K14-Cre; Pax6$^{fl/fl}$*) and age-matched controls (*Pax6$^{fl/fl}$*). **(C, C')** Pax6 staining is retained at E15.5 in CE cells of the OSE cKO mutants **(D, D')** while decreased towards presumptive conjunctiva compared to control**. (E, E')** Mosaic expression of Pax6 is observed in CE cells of OSE cKO mutants at E18.5 **(F, F'**) while downregulated at presumptive conjunctival epithelial cells. Scale bar: (**A, B,**) - 100 µm, (**A', A'', B', B''**)-50 µm **(C-F')-**50µm

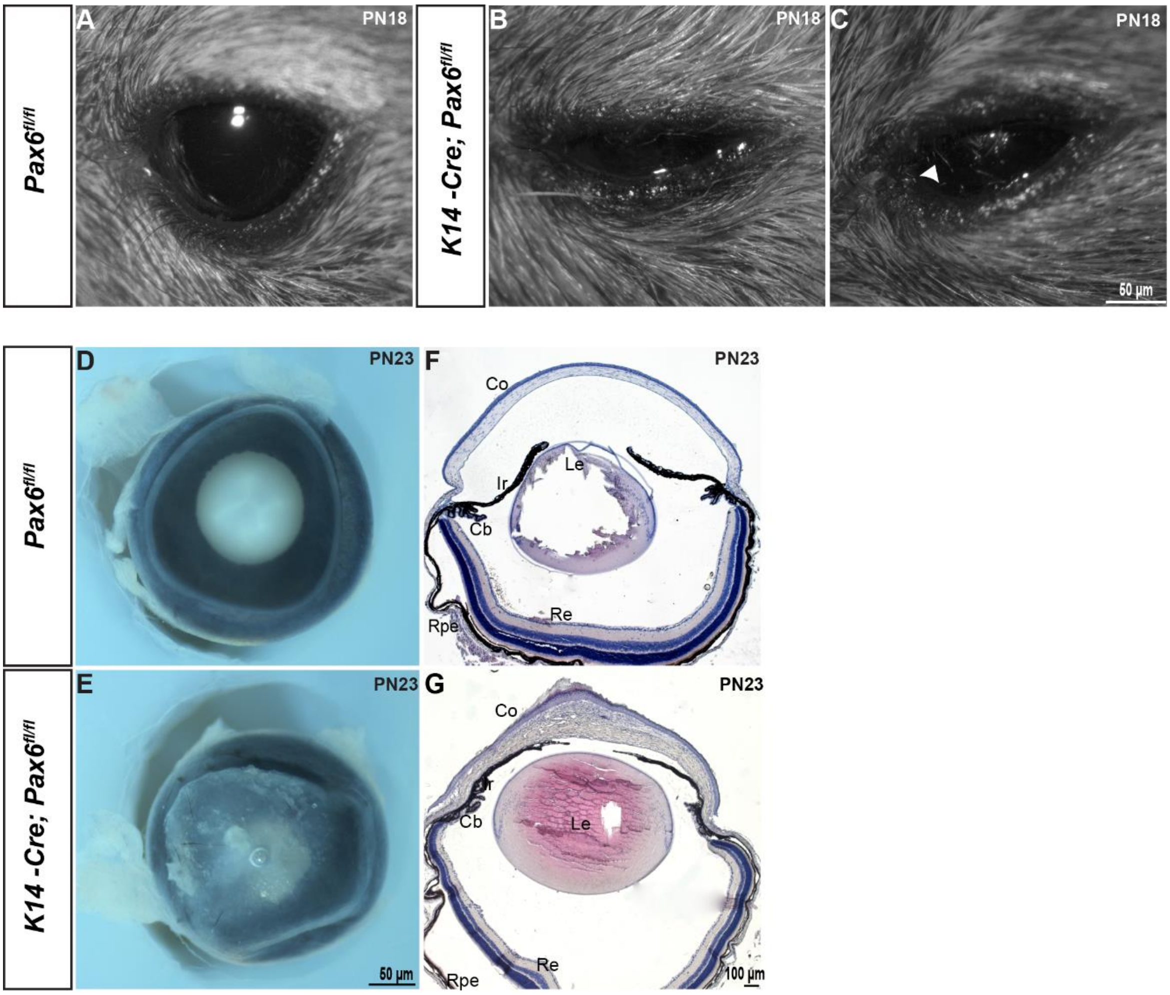


**Supplementary Figure 5: Morphological defects in OSE cKO mutants after eye-opening**

**(A)** Compared with control (*Pax6*$^{fl/fl}$) eyes, OSE cKO mutants (*K14-Cre; Pax6*$^{fl/fl}$) are with **(B, C)** smaller palpebral space and **(C)** eyelashes touching the corneal surface at PN18. **(D, E)** Control eyes at PN23 are transparent, while OSE cKO mutants have opaque eyes. **(F, G)** H&E stained coronal sections of eyes at PN23 of control and OSE cKO mutants. **(G)** Extensive thickening and keratinisation of central CE in contrast to **(F)** 4-6 stratified layer in control epithelium. **(C)** White arrowhead indicates eyelashes touching the ocular surface. Scale bar: **(A, B, C, D, E)** - 50µm **(F, G) -** 100µm

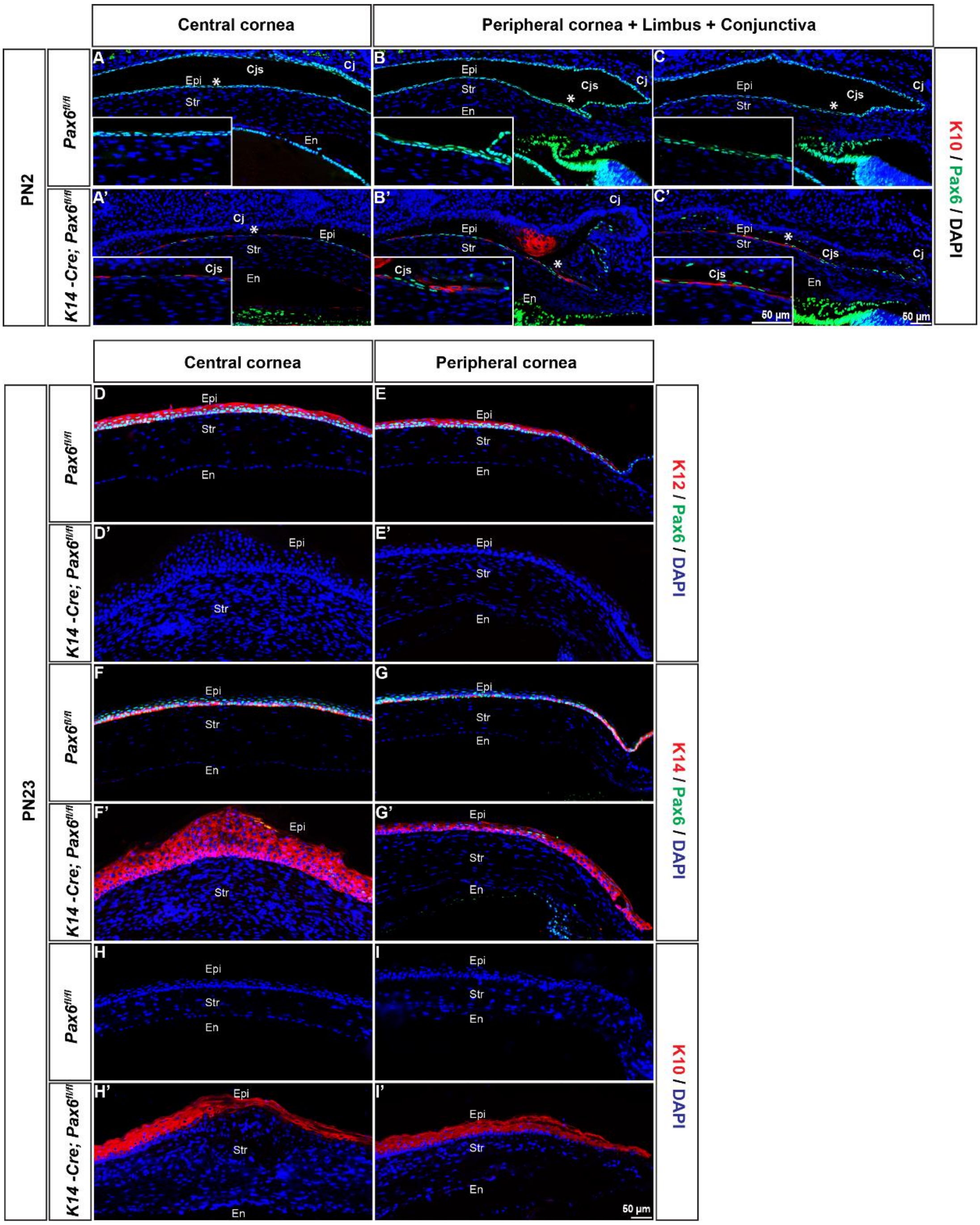


**Supplementary Figure 6: Altered keratin expression and switch in developmental fate to epidermis after eye-opening**

**(A-I')** Coronal sections of control (*Pax6$^{fl/fl}$*) and OSE cKO mutants (*K14-Cre; Pax6$^{fl/fl}$*) were stained with indicated antibodies at mentioned postnatal stages. (**A-C'**) K10$^{+}$ cells in CE and conjunctival epithelium of OSE cKO mutants at PN2. **(D-E')** K12 expression is lost in OSE cKO mutants compared to control. **(F-G')** K14 expression is expanded to all layers in OSE cKO mutants. **(H-I')** Ectopic expression of K10 in CE cells of OSE cKO mutants. Inserts in (**A-C'**) are higher magnification of (*) indicated positions. Abbreviations used: Cjs, Conjunctival sac Scale bar: **(A-I')** -50µm

| Antibody | Host | Dilution | Source |
| --- | --- | --- | --- |
| **Pax6** | Rabbit | 1:1000 | Covance (PRP-278P) |
| **K12** | Goat | 1:300 | Santa Cruz Biotec Inc, sc-17101 |
| **K14** | Mouse | 1:500 | Thermo fisher scientific, LL02,MA5-11599 |
| **K10** | Mouse | 1:500 | Thermo fisher scientific, DE-K10,MA5-13705 |
| **K4** | Mouse | 1:100 | Thermo fisher scientific, 6B10,MA1-35558 |
| **E-cadherin** | Rat | 1:300 | Thermo fisher scientific, 13-1900 |
| **β-catenin** | Rabbit | 1:1000 | Sigma, C2206 |
| **ZO-1** | Rabbit | 1:250 | Thermo fisher scientific, 61-7300 |

**Supplementary Table S1: Primary antibodies used**
